# Survey of intertidal ecosystem reveals a legacy of potentially toxic elements from industrial activity in the Skeena Estuary, British Columbia, Canada

**DOI:** 10.1101/623587

**Authors:** Tom Sizmur, Lily Campbell, Karina Dracott, Megan Jones, Nelson J. O’Driscoll, Travis Gerwing

## Abstract

Relationships between concentrations of Potentially Toxic Elements (PTEs) in estuarine sediments and their impact benthic invertebrate communities are poorly understood. We sampled and analysed PTEs in sediments and benthic invertebrates from five sites surrounding the Skeena Estuary, including sites adjacent to an abandoned cannery and a decommissioned papermill. There was no indication that sediments of the salmon cannery are polluted, but acidic sediments adjacent to the papermill contained elevated concentrations of Cd, Cr, Hg and Pb. Benthic invertebrate community assemblages confirm that sediments have recovered from prior disturbances associated with discharge of papermill sludge. Oregon pill bugs (*Gnorimosphaeroma oregonensis*), observed at all five sites, feed on the fibers associated with the papermill discharge. Thus, *G. oregonensis* are useful biomonitors for quantifying the impact of the decommissioned papermill, and similar industrial development projects, on intertidal ecosystems along the north coast of British Columbia, Canada.

## Introduction

Conflict between the economic benefits of industrial development and the potential impact of such developments on the natural environment is most clearly expressed in remote communities, entirely dependent on natural capital for survival [1–3]. Anthropogenic activities have impacted the biogeochemical cycling of Potentially Toxic Elements (PTEs) in even the most remote ecosystems on our planet [4–6]. Quantifying the disruption of natural biogeochemical cycles of these elements by anthropogenic activities is difficult if the concentrations prior to industrial development are unknown and may be redundant if investigations are only conducted after environmental damage has occurred. Therefore, it is important to conduct baseline surveys to assess the bioavailability and bioaccumulation of PTEs by organisms in remote locations to predict the likely impact of further pollution on the ecosystem.

Intertidal estuarine sediments support ecologically and economically important ecosystems that provide nursery habitats to several species (some that support important commercial fisheries), and often host a diverse and abundant fauna [7–11]. Benthic invertebrates are often used as indicators of environmental pollution since they live in sediments and are prey for many commercially important fish species [12–14]. Benthic invertebrates inhabiting estuaries are inherently resistant to physical and chemical change as they have adapted to living in a dynamic environment with wide spatial and temporal ranges of chemical and physical properties, such as pH, redox potential, salinity, and particle size [15–17]. Because the chemical and physical properties of estuarine sediment are temporally and spatially dynamic, it is difficult to predict the impact of industrial development on the fate and impact of PTEs [18]. We do not currently understand the relationship between sediment properties, food web structure and PTE bioaccumulation by sediment-dwelling invertebrates well enough to predict the impact of future emissions to the estuarine environment. The objective of this study was to explore the relationship between sediment properties and the fate of PTEs from historic industrial development in the Skeena Estuary and quantify the impact of PTEs on the diversity of benthic invertebrates inhabiting intertidal sediments.

The Skeena is British Columbia’s second largest river and provides important habitat for Pacific salmon populations, upon which Canadian First Nations communities rely [2, 8, 19]. The estuary and the intertidal areas towards the north of the Estuary along the coast of British Columbia are important nursery habitats for juvenile salmon [8, 19]. Coastal areas to the north of the estuary surrounding the small port cities of Prince Rupert and Port Edward have been extensively developed [3–5, 20]. Industrial developments include an international port, a papermill, and several canneries.

Findings of previous surveys of the benthic invertebrates inhabiting the intertidal sediment in the Skeena estuary reveal an infaunal community that is relatively undisturbed at the estuary scale, but which still shows the scars of historic disturbance at finer-grained scales [10, 15, 21, 22]. Amphipods are powerful indicator species [23–27] whose high densities throughout the Skeena estuary [10, 15] suggests that current disturbances to intertidal areas are relatively limited. Similarly, we observe 40 intertidal species in this area, including multiple species at all trophic levels within the foodweb [10, 15], and such a complex community is often associated with non-disturbed habitats [4, 28–30]. Conversely, disturbed sites are often more easily invaded by invasive species [31, 32], and Capitellidae (Capitella capitate species complex) polychaetas are often observed in disturbed habitats, particularly areas that have been organically enriched [4, 27, 28, 33]. Capitellidae polychaetas have a clumped distribution within the Skeena estuary and can be locally abundant at the scale of a 1m^2^ plot [10, 15]. Similarly, abundant but localized populations of the invasive Cumacea *Nippoleucon hinumensis*, have been observed in the Skeena estuary [10, 15], and on other historically impacted mudflats along BC’s north coast [27]. While the universal presence and high abundances of amphipods, coupled with the complexity of the intertidal community at the estuary scale strongly suggest that these intertidal communities are relatively undisturbed, the locally abundant populations of invasive species and Capitellidae polychaetas offer contradictory evidence. These indicators of disturbance suggest that either current small-scale disturbances are occurring, or this biological signal is a remnant of past disturbances.

Persistent localised biological signals of historic disturbances in mudflats that are currently relatively undisturbed have been observed before on the north coast of BC [27]. To the best of our knowledge, no current disturbances are occurring in the Skeena estuary that can explain the small-scale indicators of disturbance observed in the biological community [10, 15, 22, 27]. Therefore, we hypothesise that fine-grained biological indicators of disturbance in the Skeena estuary are a legacy of past disturbances [10, 15]. It is likely that these past disturbances are at least partially related to discharge from the papermill, which was released into the immediate near shore area (Porpoise Bay), strongly depressing the invertebrate communities in this area during the 1970s [20, 34, 35]. As such, we postulate that these intertidal areas have been passively recovering from this disturbance for ∼50 years.

Considerable developments have been proposed in the Skeena estuary, including oil and gas pipelines, super-tanker routes, potash loading facilities, and a liquid natural gas (LNG) terminal. The Skeena salmon run contributes an estimated $110 million dollars annually to the local economy [36], so pollution of the intertidal nursery habitat of the Skeena Estuary could have devastating consequences on both the economy and ecosystem [37, 38]. It is therefore critical to predict the impact that future developments will have on the bioaccumulation of PTEs by benthic invertebrates inhabiting intertidal sediments of the Skeena Estuary. This work presents a survey of intertidal sediments and invertebrates at five locations north of the Skeena estuary to examine baseline levels of PTE concentrations at sites that may be recovering from previous disturbances and identify organisms that could be used to biomonitor the impact of future industrial developments.

## Methods

### Field sampling

Sediment cores and benthic invertebrates were collected from five locations north of the Skeena Estuary (Fig 1) during July 2017: Cassiar Cannery (CC), Inverness Passage (IP), Tyee Banks (TB), Papermill Bay (PB), and Wolfe Cove (WC). Cassiar Cannery (N54° 10’ 40.4, W130° 10’ 40.4) is a former salmon cannery that closed in 1983 and is now an ecotourism lodge. Inverness passage (N54° 10’ 05.9, W130° 09’ 40.4), a mudflat ∼3 km closer to the Skeena mouth than Cassiar Cannery, has intertidal habitat similar to Cassiar Cannery, but was never directly impacted by a cannery or other anthropogenic development. Tyee Banks (N54° 11’ 59.1, W129° 57’ 36.7) is located 20km upstream from the mouth of the Skeena River on a large intertidal mudflat. At some point in the past, this area had a small-scale sawmill operating and accumulations of sawdust and woodchips are still present in the upper intertidal sediment. Papermill Bay (N54° 13’ 59.3, W130° 17’ 07.5) is a small bay located directly adjacent to a large decommissioned papermill. Finally, Wolfe Cove (N54° 14’ 33.0, W130° 17’ 34.5) is a mudflat located approximately 1 km from the papermill. The papermill was closed in 2001, ceasing all operations and discharge [39–41].

**Fig 1.**
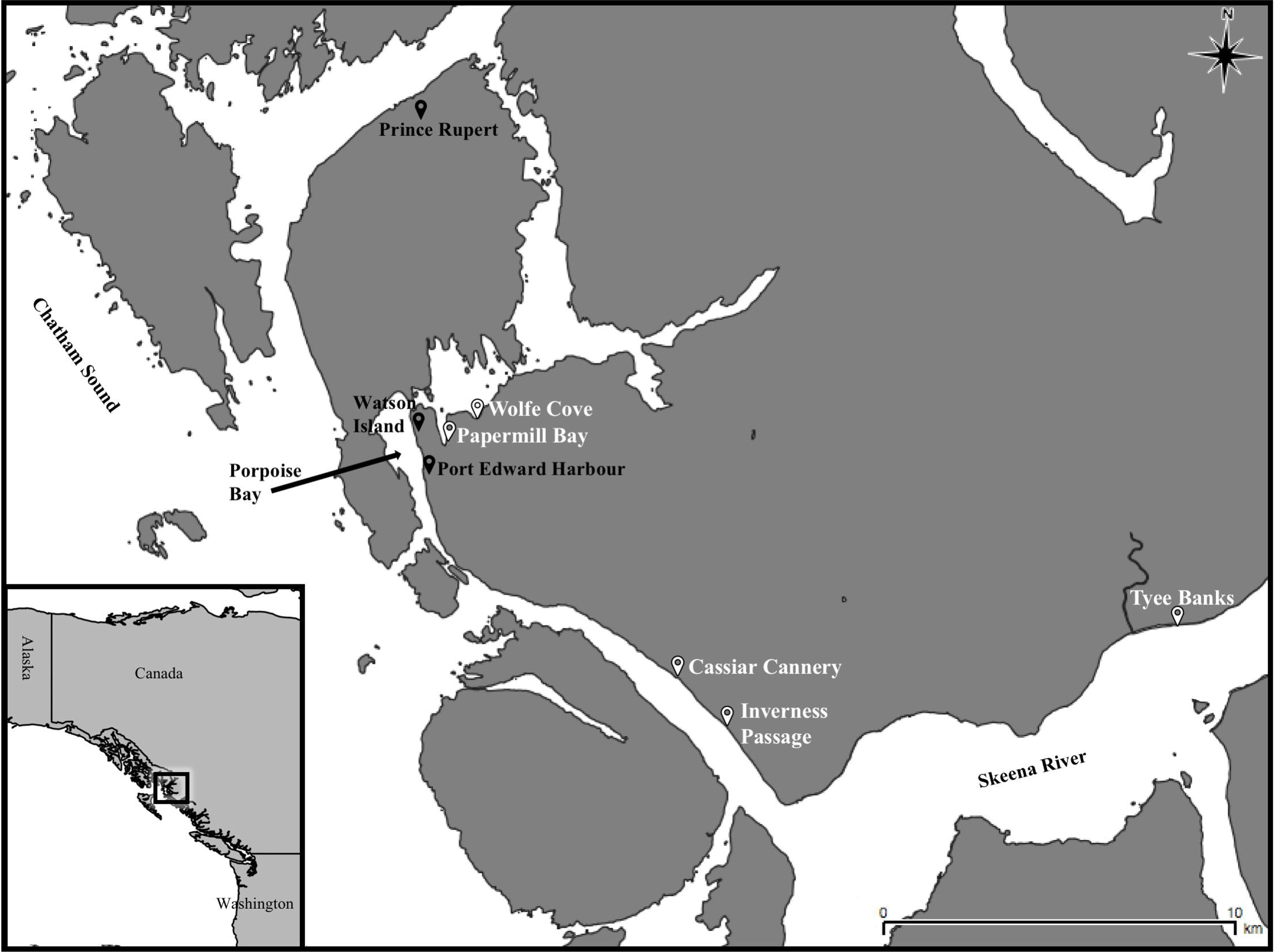
Map showing sites adjacent to the Skeena Estuary, British Columbia, Canada where intertidal mudflats at Papermill Bay, Wolfe Cove, Cassiar Cannery, Inverness Passage, and Tyee Banks were sampled during this project.

Sediments were sampled by pushing polycarbonate cores (5 cm diameter, 20 cm length) into the mud with a rubber mallet, digging out with a spade, and wrapping with cling film, before extruding and dividing into strata; 0-5 cm, 5-10 cm, 10-15 cm and 15-20cm. Sediment samples from each strata were dried at 40 °C for 48 hours and shipped to the UK for analysis. A total of 15 cores were collected from each site along five transects comprising three samples taken from the upper, mid and lower shore in a random stratified sampling design [42]. Sediments from the upper, mid and lower shore (but the same strata) were pooled and homogenized prior to analysis. At each location that a sediment core was collected, two more cores (7 cm diameter) were taken to a depth of at least 5 cm (well into the anoxic layer), transferred to a plastic bag in the field and then passed through a 250µm stainless steel sieve to retain sediment-dwelling benthic infauna. This infauna sampling strategy was supplemented by opportunistic digging in areas of the site in which infauna were clearly present (e.g. burrow openings or surface casting) to collect macrofauna. All invertebrates found in sufficient quantities for chemical analysis were identified to the species level [10, 15], pooled into a single sample for each site, rinsed in deionised water, frozen, shipped to the UK, and freeze-dried prior to analysis.

### Laboratory analysis

The particle size distribution of sediments was determined using laser granulometry (Malvern Mastersizer 3000). Sub-samples were then ground to a fine powder using an agate ball mill and analysed for total organic carbon and nitrogen content using a Thermo Scientific Flash 2000 Organic Elemental Analyser. Details of quality control can be found in the supporting information file. Sediment pH was determined in a soil-water suspension after shaking with water for 15 min at a 1:10 w/v ratio based on BS7755-3.2 [43]. The total concentration of PTEs in sediments was determined by ICP-MS (Inductively Coupled Plasma Mass Spectrometry) analysis of 0.5 g sediment samples digested in reverse aqua regia (9 ml of nitric acid and 3 ml of hydrochloric acid) using a MARS 6 microwave digestion system, based on EPA [44]. After preliminary analysis of a large range of PTE concentrations, the elements selected for further investigation were Cadmium (Cd), Cobalt (Co), Chromium (Cr), Nickel (Ni), Lead (Pb) and Zinc (Zn). Details of quality control can be found in the supporting information file. The total concentration of mercury in sediments was determined using thermal degradation – gold amalgamation atomic absorbance spectroscopy as outlined in EPA [45] using a Nippon MA-3000 analyser. Details of quality control can be found in the supporting information file. The bioavailability of PTEs to biota was estimated by extracting 2.5 g of sediments with 25 ml of 0.05M EDTA (Ethylenediaminetetraacetic acid) at 20°C for one hour, centrifuging, filtering and analysing PTEs (As, Cd, Co, Cr, Cu, Ni, Pb and Zn) in the extract using ICP-OES (Inductively Coupled Plasma Optical Emission Spectrometry). The total concentrations of PTEs in invertebrates were determined by ICP-MS analysis of 0.5 g of sample digested in slightly diluted nitric acid (2 ml of ultra-pure water and 8 ml of nitric acid) using a MARS 6 microwave digestion system.

### Statistical analysis

The influence of site and sediment depth on sediment PTE concentrations and properties was quantified using analysis of variance (ANOVA) and permutational multivariate analysis of covariance (PERMANCOVA) [46, 47]. The relationship between PTE concentrations and sediment properties was first tested using PRIMER’s RELATE function [48]. This function compares two resemblance matrices looking for any relationships. Relationships were further explored using principal component analyses (PCA) on the variance-covariance matrix of all sediment PTE and sediment property data. Relationships between the PTE concentrations observed in the sediments and in the collected invertebrates were examined in several configurations using RELATE and plotted using non-metric multidimensional (nMDS) scaling plots. A full description of the statistical analysis undertaken is provided in the supporting information file and outputs provided in S1, S2 and S3 Tables.

## Results and Discussion

### Inverness Passage, Wolfe Cove and Tyee Banks are suitable reference sites

The PERMANCOVA analysis indicates that properties of sediments (pH, median particle diameter, C and N) are significantly influenced by both site and sediment depth, with site explaining more than 50% of the observed variance in the data (S1 Table). There is no significant (p > 0.05) difference in the median sediment particle diameter between Inverness Passage and Cassiar Cannery, or between Wolfe Cove and Papermill Bay (Fig 2 and S3 Table). This observation supports our assumption that the sediment deposited in the sites potentially contaminated by industrial emissions (Cassiar Cannery and Papermill Bay) and their respective proximal reference sites (Inverness Passage and Wolfe Cove) have the same geogenic origin and are subject to a similar depositional environment. Thus, any differences in sediment chemical properties between potentially contaminated sites and their proximal reference sites we infer is due to anthropogenic influences.

**Fig 2.**
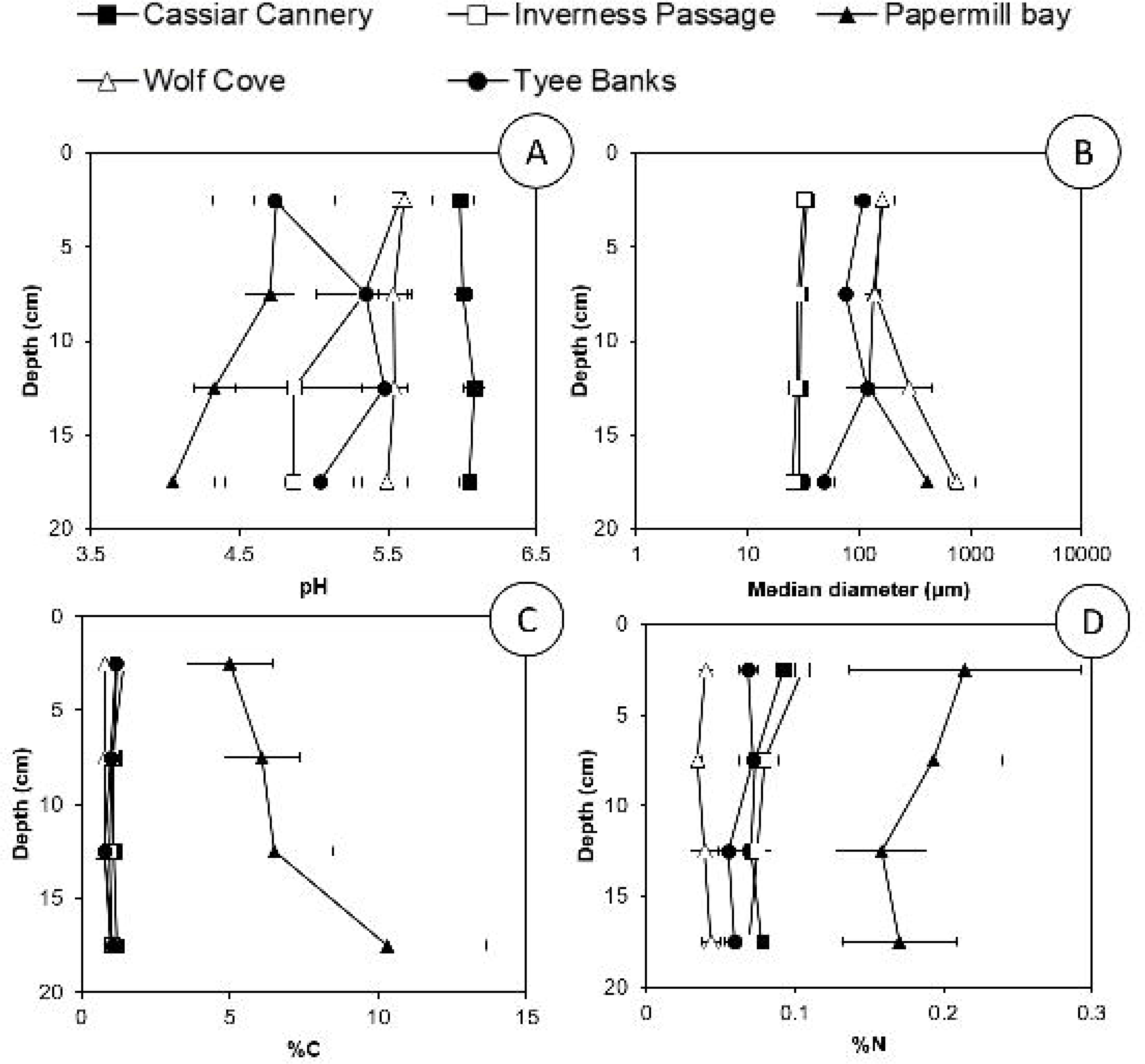
Sediment properties; Sediment pH (A); Median sediment particle diameter (B); Percentage Organic Carbon content (C); and Percentage Total Nitrogen content (D) presented at four depths in sediments sampled from Cassiar Cannery, Inverness Passage, Papermill Bay, Wolfe Cove and Tyee Banks intertidal mudflats.

Tyee Banks has a significantly (p < 0.05) larger median particle size than Inverness Passage and Cassiar Cannery and a significantly (p < 0.05) smaller median particle size than Wolfe Cove and Papermill Bay (Fig 2 and S3 Table). The location of Tyee Banks is closer to the mouth of the Skeena River and, therefore, only receives sediment from the Skeena, whereas the other four sites may also receive sediment from the Nass River, to the north of the Skeena Estuary [49]. However, because Tyee Banks is much further from potential emission sources, the site can be used to determine whether proximal reference (Inverness Passage and Wolfe Cove) sites are contaminated with PTEs from the adjacent industrial activity.

### Papermill Bay sediments are contaminated with Cd, Cr, Hg and Pb

The PERMANCOVA analysis (S2 Table) indicates significant differences in sediment total and EDTA extractable PTE concentrations between different sites, but not between different depths. We observed significantly (p < 0.05) greater concentrations of As, Cr, Hg and Pb in the Papermill Bay sediments than the proximal reference sediments at Wolfe Cove (Fig 3 and S3 Table). However, Co, Cu and Ni concentrations were significantly (p < 0.05) lower in the Papermill Bay sediments (Fig 3 and S3 Table). A Geoaccumulation index, calculated following [50], using Wolfe Cove as an uncontaminated reference site, indicates that Papermill Bay is ‘unpolluted’ with As, Cd, Co, Cr, Cu, Ni and Zn, but ‘unpolluted to moderately polluted’ with Pb and ‘moderately polluted’ with Hg (Table 1). When Tyee Banks is used as the reference site, Papermill Bay is classified as ‘unpolluted to moderately polluted’ with Cd, Cr and Hg. Thus, there is evidence to suggest that the sediments in Papermill Bay are contaminated with Cd, Cr, Hg and Pb, most likely emanating from the sludge [51] discharged by the decommissioned papermill on Watson Island (Fig 1).

**Table 1.** Geoaccumulation index (Igeo)^a^ for potentially toxic elements in sediments from Cassiar Cannery (CC) and Papermill Bay (PB) using reference sites Tyee Banks (TB), Wolfe Cove (WC) and Inverness Passage (IP).

| Site | Reference site | As | Cd | Co | Cr | Cu | Hg | Ni | Pb | Zn |
| --- | --- | --- | --- | --- | --- | --- | --- | --- | --- | --- |
| CC | IP | -0.99 | -0.71 | -0.78 | -0.57 | -0.69 | -0.69 | -0.67 | -0.58 | -0.63 |
|  | TB | -0.48 | -0.33 | -0.44 | -0.27 | -0.01 | -0.13 | -0.26 | 0.08 | -0.24 |
| PB | WC | -0.10 | -0.25 | -1.08 | -0.32 | -1.14 | 1.61 | -0.82 | 0.74 | -0.60 |
|  | TB | -1.51 | 0.39 | -0.90 | 0.35 | -0.19 | 0.20 | -1.01 | -0.02 | -0.45 |

<sup>a</sup> Geoaccumulation index is calculated as $I_{geo} = \log_2 \frac{C_n}{1.5 \times B_n}$
when $C_n$ = Concentration measured in sediment at site and $B_n$ = Concentration measured in sediment at reference (background) site.
Sites are classified as unpolluted if $I_{geo} \leq 0$ , unpolluted to moderately polluted if $0 < I_{geo} \leq 1$ , moderately polluted if $1 < I_{geo} \leq 2$ , moderate to strongly polluted if $2 < I_{geo} \leq 3$ , strongly polluted if $3 < I_{geo} \leq 4$ , strongly to extremely polluted if $4 < I_{geo} \leq 5$ , and extremely polluted if $I_{geo} > 5$ .

**Fig 3.**
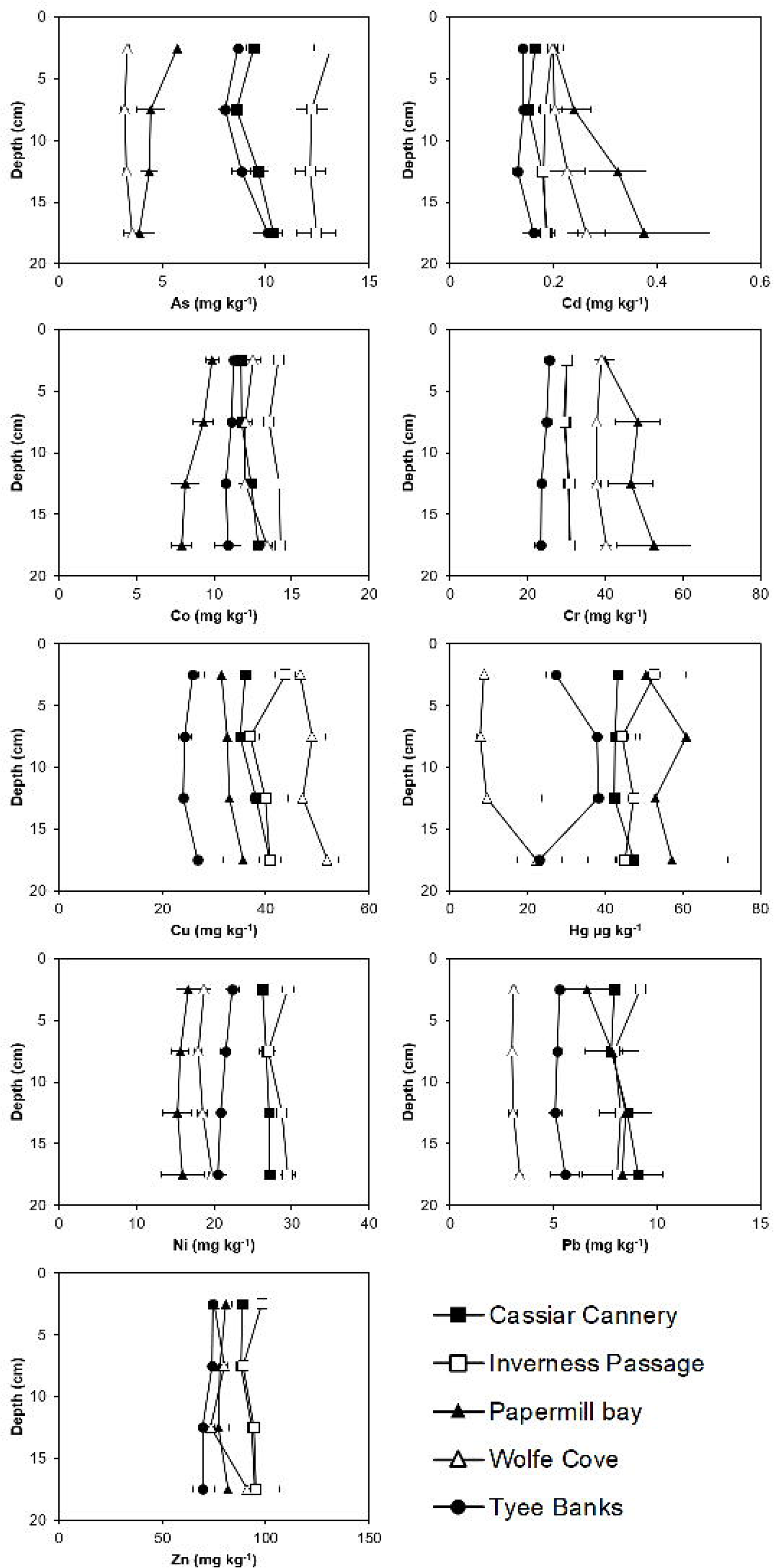
Total concentrations of Potentially Toxic Elements (As, Cd, Co, Cr, Cu, Hg, Ni, Pb and Zn) with depth in sediments sampled from Cassiar Cannery, Inverness Passage, Papermill Bay, Wolfe Cove and Tyee Banks intertidal mudflats

Discharges by papermills have previously been associated with pollution of the environment with PTEs, including Cr, Cd, Cu, Pb and Zn [52, 53]. This pollution associated with papermill operations can originate from a number of sources, such as metals in the wood entering the papermill, or atmospheric deposition of metals from the smoke stacks [52]. PTE-containing materials may also be used during the pulp making processes. For example, chromate bricks are used in the recovery furnace of kraft paper mills to reduce chemical attack by the spent liquors [54]. However, Cr concentrations observed in the sediments of Papermill Bay (Fig 3) are considerably lower than the 52.4 mg kg^−1^ observed in a deposit of pulp covering the sediments of a Swiss lake [55], or concentrations of up to 197 mg kg^−1^ observed in a fibrous deposit in the Ångermanälven river estuary, on the east coast of Sweden (Apler et al 2019).

The carbon and nitrogen content of the sediments at Papermill Bay were significantly (p < 0.05) and considerably greater than all of the other sites (Fig 2 and S3 Table). It was evident from visual inspection of the sediments themselves that they contained a high proportion of organic fibers, presumably discharged into the bay from the papermill during the operational phase. The PTEs observed to be elevated in the Papermill Bay sediments (Cd, Cr, Hg and Pb) all bind strongly to organic matter in sediments [56–58] and so their presence could be due to (i) the discharge and deposition of PTE contaminated organic material, (ii) the discharge of organic material alongside the discharge of PTEs in contaminated effluent that later bind with the organic material, or (iii) the discharge of organic material which binds to naturally present PTEs in the sediment and prevents their export to the overlying water.

In contrast to the observations made in the Papermill Bay sediments, none of the PTEs analysed in the Cassiar Cannery sediments were found to be present in significantly (p > 0.05) greater concentrations than the Inverness Passage sediments (Fig 3 and S3 Table). In fact, Co and Ni concentrations were significantly lower in Cassiar Cannery sediments than at Inverness Passage. When Inverness Passage is used as a reference site, all the PTEs analysed can be classified by the Geoaccumulation index as ‘unpolluted’ (Table 1). This observation is a clear indicator that any impact resulting from either the former use of the Cassiar Cannery site (up until 1983) as a salmon cannery, or its current use as an ecotourism lodge, can no longer be observed in the concentrations of PTEs in the top 20 cm of sediment at this site. Any sediment contaminated by cannery waste at this site may have been buried by fresh sediment, rendering the PTEs inaccessible to benthic invertebrates.

### Relationship between sediment properties and total and labile PTE concentrations

Alongside greater total concentrations of Cd, Cr, Hg and Pb in the Papermill Bay sediments, compared to Wolfe Cove, we observed significantly (p < 0.05) greater EDTA extractable concentrations of Cd, Co, Cr, Ni, Pb and Zn (Fig 4 and S3 Table), indicating a greater availability of these elements to benthic invertebrates [59]. The significantly (p < 0.05) greater carbon content and the slightly finer texture of the Papermill Bay sediments, compared to Wolfe Cove (Fig 2 and S3 Table) should provide more sites for PTEs to bind to [58, 60, 61]. Thus, the greater concentrations of EDTA extractable PTEs is largely due to the Papermill Bay sediments being significantly (p < 0.05) more acidic than the Wolfe Cove sediments. Lower pH results in greater competition between PTEs and hydrogen ions for the binding sites on the sediment surface and leads to greater PTE lability [62, 63].

**Fig 4.**
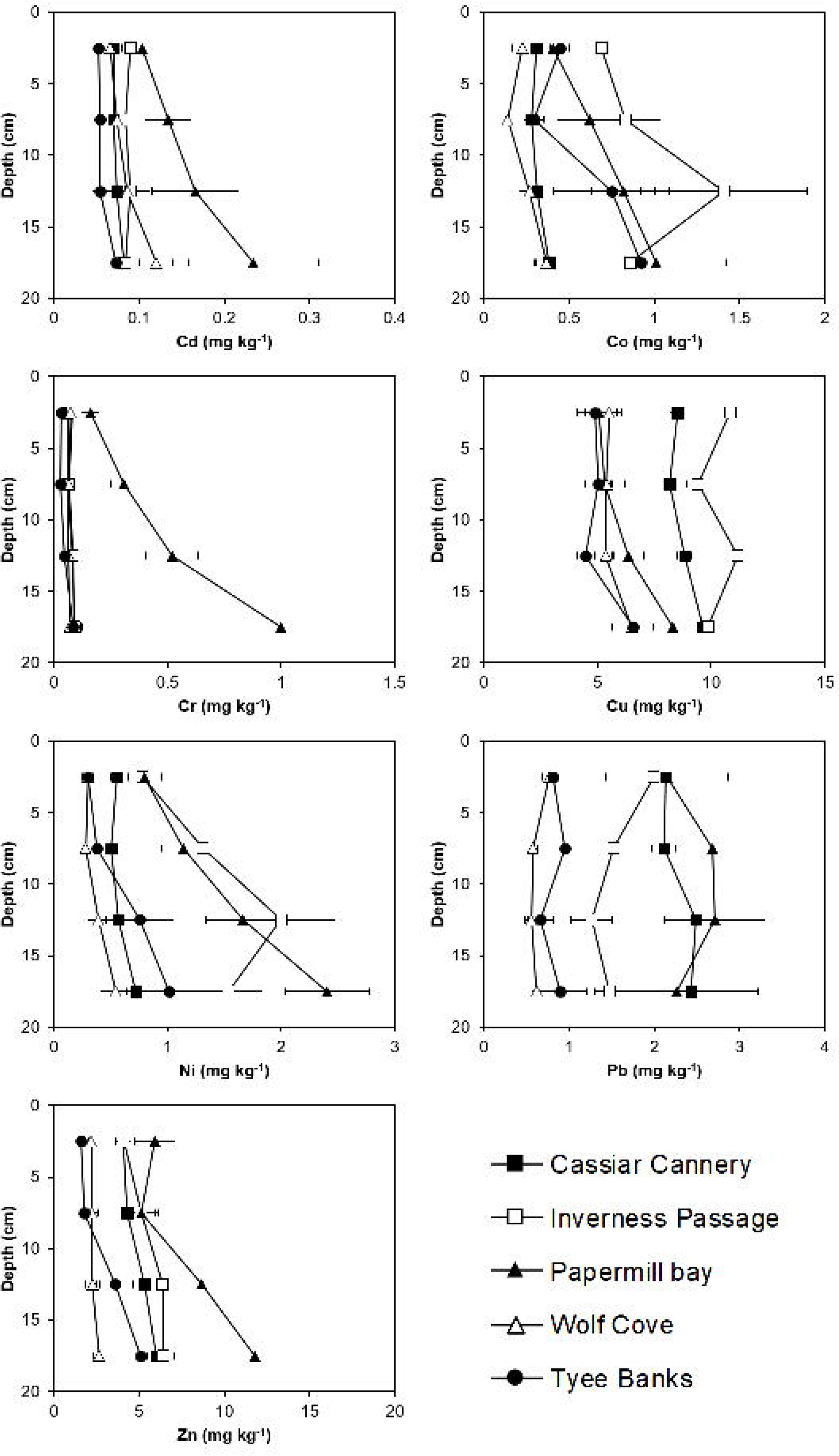
EDTA extractable (potentially bioavailable) Potentially Toxic Elements (Cd, Co, Cr, Cu, Ni, Pb and Zn) with depth in sediments sampled from Cassiar Cannery, Inverness Passage, Papermill Bay, Wolf Cove, and Tyee Banks intertidal mudflats

The RELATE revealed a statistically significant correlation between the concentration of potentially toxic elements and the sediment physicochemical properties resemblance matrices (Rho: 0.46; *p* = 0.001, Permutations = 9999). The relationship between total and labile PTE concentrations and sediment properties was further explored using PCA (Fig 5). The principal component scores plot (Fig 5B) reveals a clear separation between the Papermill Bay sediments and the proximal and remote references sites of Wolfe Cove and Tyee Banks, but no separation between Cassiar Cannery and Inverness Passage, re-enforcing the conclusion that the sediments of Cassiar Cannery show little evidence of anthropogenic contamination. Principal component 2 (Fig 5), separates the Inverness Passage and Cassiar Cannery sediments from the other three sites (Papermill Bay, Wolfe Cove, and Tyee Banks), which is attributed to a different geochemical matrix composition at Cassiar Cannery and Inverness Passage, as also observed by DelValls et al. [64]. Principal component 1 (Fig 5) reveals a separation with depth in the Papermill Bay sediments. Papermill Bay sediments have greater total and EDTA extractable concentrations of several PTEs, higher C and N, and lower pH, all of which also increase with depth in the Papermill Bay sediments. This observation of deeper layers of sediment with a lower pH and higher availability of PTEs indicates that contaminated sediment is overlain by less contaminated sediment, deposited since discharge of sludge ceased. Although most benthic invertebrates live in surficial sediments, it is important to consider the likelihood that deeper sediment will be exposed and mobilized either by development projects [65], or by bioturbation from deep burrowing invertebrates [66, 67].

**Fig 5.**
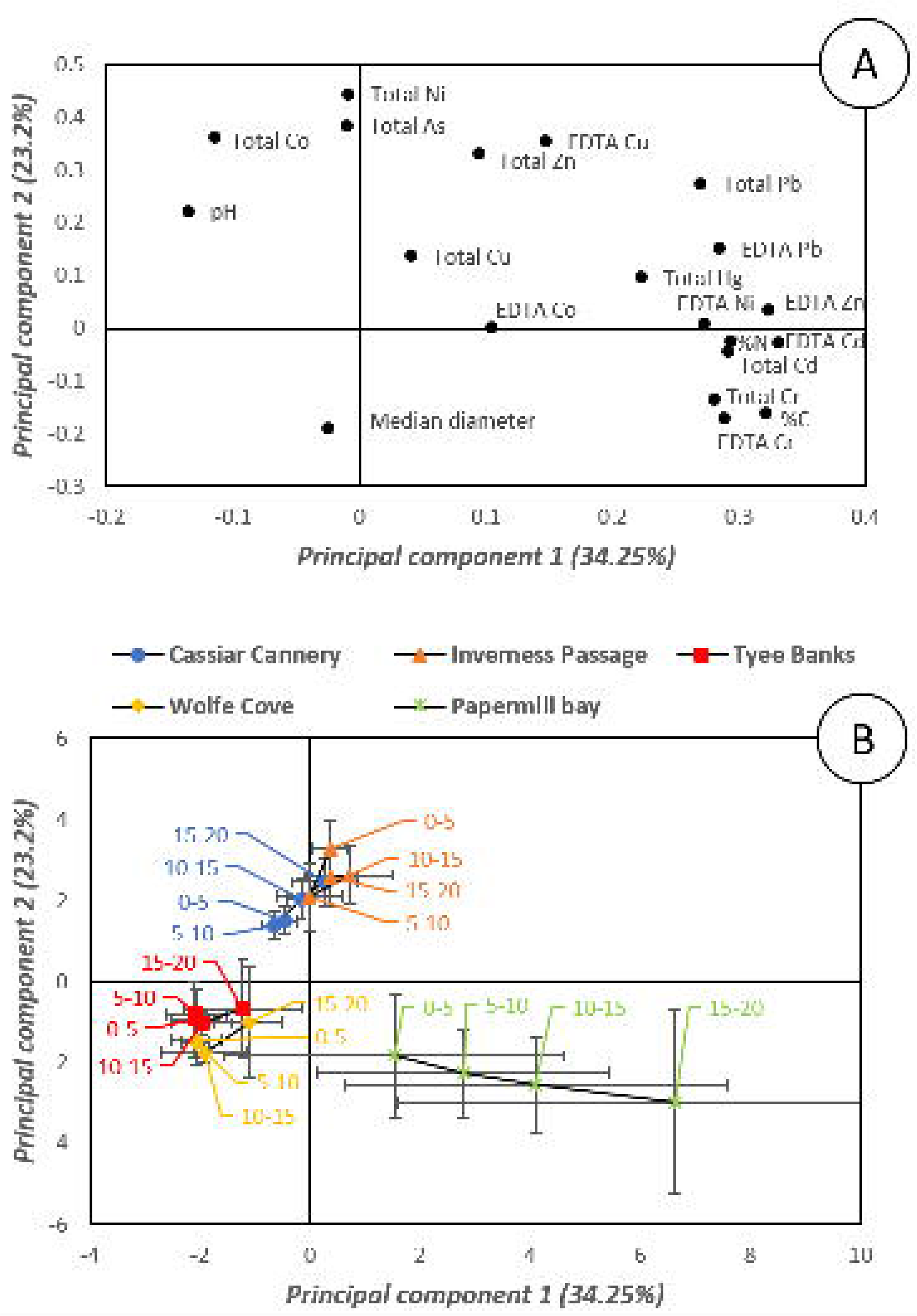
Principal Components Analysis of sediment properties (pH median sediment particle diameter, Percentage Organic Carbon content and Percentage Total Nitrogen content) and total and EDTA extractable Potentially Toxic Elements (As, Cd, Co, Cr, Cu, Hg, Ni, Pb and Zn) in sediments sampled 0-5 cm, 5-10 cm, 10-15 cm and 15-20cm from Cassiar Cannery, Inverness Passage, Papermill Bay, Wolf Cove and Tyee Banks intertidal mudflats. (A) is the latent vectors for each variable plotted in the plane of principal component one and principal component two, (B) is the principal component scores of all the samples plotted in the plane of principal component one and principal component two. Error bars represent the standard deviations of 5 replicate samples taken from 5 different transects.

### Sediments host a healthy invertebrate population with no relationship between PTE concentrations in sediments and invertebrates

Overall, 22 different taxa (listed in S4 Table, along with PTE concentrations) were observed in this study, with Wolf Cove and Cassiar Cannery the most biodiverse sites, with 16 and 14 species collected, respectively. These taxa represent a subset of the 40 intertidal species that are commonly observed in the Skeena Estuary [10, 15]. Findings of previous studies in the Skeena Estuary reveal an infaunal community predominantly dominated by cumaceans, polychaetes, oligochaetes, nematodes, copepods, amphipods, and bivalves [15, 68]. The 40 intertidal species observed in this area include multiple species at all trophic levels within the food web [10, 15], and such a complex community is often associated with non-disturbed habitats [4, 28]. This complexity was also observed in our study, as *Alitta brandti, Neries vexillosa, Glycinde picta,* ribbon worms and crabs can act as predators, while *Macoma balthica*, isopods, amphipods, and sessile polychaete worms are primary consumers [47, 69, 70]. We therefore provide evidence to support our previous research that indicates that the intertidal ecosystem has been passively recovering for ∼50 years from past disturbances related to discharge from the papermill, which was released into the immediate near shore area (Porpoise Bay), strongly depressing the invertebrate communities in this area during the 1970s [20, 34, 35] and currently exhibit a community that is relatively healthy.

When all invertebrates collected at all five sites are considered, we found no relationship between PTE concentrations in invertebrates and either total (Rho: 0.27; *p* = 0.32, Permutations = 9999), EDTA extractable (Rho: 0.24; *p* = 0.79, Permutations = 9999), or both total and EDTA extractable (Rho: 0.78; *p* = 0.08, Permutations = 9999) PTEs in the sediment. This finding is confirmed by inspecting a nMDS plot of invertebrate PTE loadings, which includes all organisms collected at all five sites (Fig 6A). Bivalves of the same species (e.g. *M. balthica, Mytilus edulis*, and *Mya arenaria*), collected at different sites, seem to cluster in multidimensional space. There is slightly more separation in the PTE concentrations measured in polychaete worms (e.g. *Abarenicola pacifica, N. vexillosa, Paranemertes peregrina, G. picta, A. brandti, Nephtys caeca, Neotrypaea californiensis*, and *Streblospio benedicti*) which are the cause of much of the dissimilarity in the dataset.

**Fig 6.**
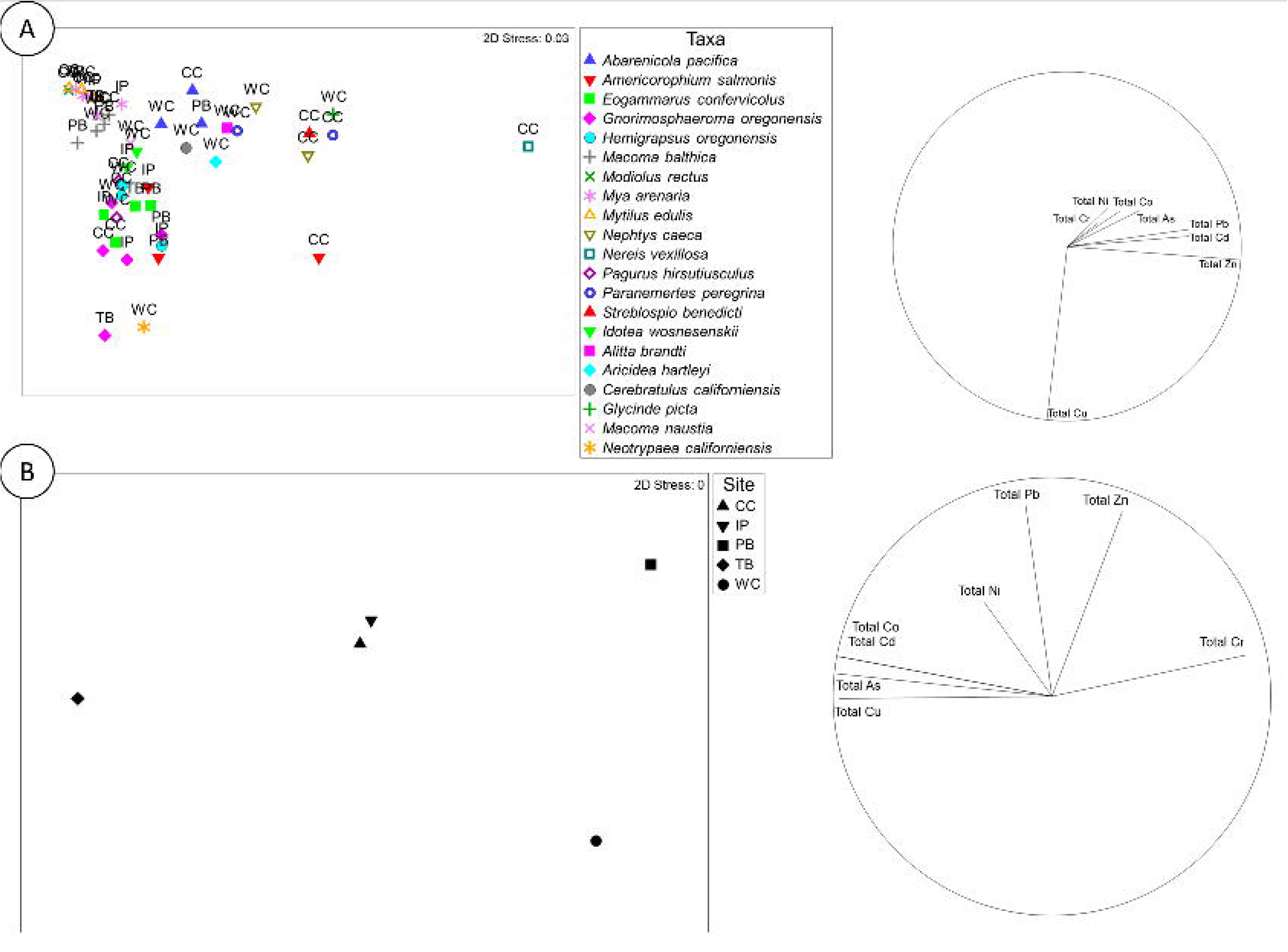
Non-metric multidimensional scaling plots (nMDS) of invertebrate PTE concentrations and the vector overlay (left hand side) of five intertidal mudflats along the north coast of British Columbia, Canada. CC: Cassiar Cannery. WC: Wolfe Cove. IP: Inverness Passage. PB: Papermill Bay. TB: Tyee Banks with (A) the entire dataset considered or (B) only including the two benthic invertebrates found at all five sites; Baltic clams (*Macoma balthica*) and Oregon pill bugs (*Gnorimosphaeroma oregonensis*).

### The suitability of *G. oregonensis* as biomonitors of sediment PTE bioavailability

The selection of one or more of benthic invertebrate as a biomonitor of PTE pollution needs to consider how cosmopolitan their distribution is and the pathway by which they are exposed to the PTE [71]. Mussels and clams acquire food from the water column by suspension feeding, and clams are also able to acquire food from surface sediments by way of their siphon. Isopods are more motile and acquire food by ingesting organic debris, usually on the surface of the sediment. Polychaete worms, as a class, have a more diverse range of feeding strategies, including suspension feeding with tentacles or a mucus net, deposit feeding selectively at the surface or at depth, and predation of other infauna [72, 73]. This large diversity of feeding strategies may have led to the greater range of PTE concentrations observed in Fig 6A, compared to the bivalves and crustacea. Furthermore, the soft tissues of polychaete worms may only represent exposure to PTEs in the recent past due to large temporal fluctuations [74–76]. The whole body (including soft tissues and shells) of bivalves may better represent a record of time-averaged bioavailability of PTEs in the water column over the lifetime of the organism, albeit not without difficulties in interpretation [77, 78].

There were only two benthic invertebrate species observed at all five sites; Baltic clams (*M. balthica*) and Oregon pill bugs (*G. oregonensis*). When PTE concentrations in *M. balthica*, and *G. oregonensis* was compared to sediment concentrations, no relationship was observed with either total (Rho: 0.31; *p* = 0.19, Permutations = 9999), EDTA extractable (Rho: −0.0.; *p* = 0.48, Permutations = 9999), or both total and EDTA extractable (Rho: 0.21; *p* = 0.28, Permutations = 9999) sediment PTE concentrations. These findings contrast to numerous articles in the literature quantifying relationships between sediment and benthic invertebrate PTE concentrations [58, 71, 79, 80]. When we plot the PTE concentrations in *M. balthica*, and *G. oregonensis* in multidimensional space (Fig 6B), we reveal that the elemental profiles of *M. balthica*, and *G. oregonensis* contrast greatly. *M. Bathica* collected at Papermill Bay contained lower concentrations of Cr, Co, Ni, Zn, Cd, As and Pb than those collected at the other four sites, but higher concentrations of Cu. In contrast, *G. oregonensis* collected at Papermill Bay have higher concentrations of Cr, Ni, Zn, and Pb than those collected at the other four sites.

Biota-Sediment Accumulation Factors (BSAFs) were calculated by dividing the concentrations of PTEs in the invertebrate tissues by the average concentration in the sediments from the site from which the invertebrates were collected (average across all transects and depths). Relationships between sediment properties and BSAFs reveal the importance of pH in explaining the difference in the bioaccumulation of PTEs by *M. balthica*, and *G. oregonensis*. For most PTEs there is a positive relationship between pH and the BSAF for *M Balthica* (Table 2), including a significant relationship with the Cr BSAF (Fig 7A). This is an unexpected finding that does not have an immediately obvious explanation. However, we observe a negative relationship between sediment pH and the BAF of most PTEs for *G. oregonensis* (Table 2), including Cr (Fig 7B). This relationship is intuitive since metal cations dissociate from mineral surfaces at lower pH levels [62, 63]. Because *G. oregonensis* feed on organic material on the surface of the sediments, the concentrations of PTEs associated with acidic sediments contaminated by papermill sludge at Papermill Bay, are more likely to be assimilated by *G. oregonensis* and become bioaccumulated in their tissues. A survey of intertidal areas adjacent to papermills on the British Columbia coastline (including the papermill on Watson island) identified *G. oregonensis* as tolerant of papermill impacted shorelines [81]. The greater abundance of isopods (including *G. oregonensis*) in close proximity to British Columbian papermills, also observed by Robin et al. [82], is attributed to the provision of pulp fibers from the mill as a food source creating a stressed environment which enables these isopods to thrive.

**Table 2.** Correlation coefficients for the relationship between sediment pH and Biota-Sediment Accumulation Factors of 8 potentially toxic elements in Baltic clams (*Macoma balthica*) and Oregon pill bugs (*Gnorimosphaeroma oregonensis*) at all five sampling locations in the Skeena Estuary.

| Species | As | Cd | Co | Cr | Cu | Ni | Pb | Zn |
| --- | --- | --- | --- | --- | --- | --- | --- | --- |
| *M Balthica* | 0.4397 | 0.2564 | 0.9162 | 0.8255* | -0.8177 | 0.9461 | 0.5788 | 0.896 |
| *G. oregonensis* | -0.1744 | 0.3758 | -0.7362* | -0.8688* | -0.1895 | -0.8117 | -0.2083 | -0.7732* |

\* = $p < 0.05$

**Fig 7.**
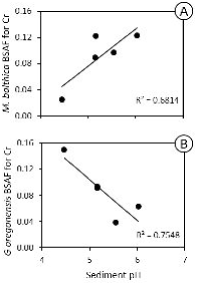
The relationship between Chromium (Cr) Biota-Sediment Accumulation Factor (BSAF) and sediment pH for (A) Baltic clams (*Macoma balthica*) and (B) Oregon pill bugs (*Gnorimosphaeroma oregonensis*) sampled at five intertidal mudflats along the north coast of British Columbia, Canada.

## Conclusions

We found evidence to indicate that the discharge of papermill sludge from the decommissioned papermill on Watson Island has changed the sediment geochemistry at Papermill Bay by reducing the pH, and increasing the total and EDTA extractable concentrations of Cd, Cr, and Pb. However, the benthic invertebrate community composition confirms that the population has recovered from previous disturbance. Oregon pill bugs (*G. oregonensis*) were one of only two benthic invertebrate species observed at all five sites we visited. *G. oregonensis* are a cosmopolitan species along the north coast of British Columbia and are tolerant of sites contaminated with papermill effluent because they use the fibers discharged as a food source. Thus, we conclude here that *G. oregonensis* make an excellent candidate biomonitor species to assess recovery from the environmental impact of the papermill on Watson Island and monitor the future impacts of similar industrial developments in the region.

## Acknowledgements

Sean Allen is acknowledged for fieldwork assistance. Ilse Kamerling is acknowledged for assisting C and N analysis. Anne Dudley is acknowledged for assisting ICP-OES analysis. Andy Dodson is acknowledged for assisting ICP-MS analysis. Accommodation and access to the shoreline were graciously provided by Cassiar Cannery Ecotourism lodge.

## Supporting information captions

The supporting information file contains additional text that elaborates on the quality control and statistical analysis undertaken in addition to the following tables:

**S1 Table. PERMANCOVA showing sediment properties (pH, median particle diameter, C and N) varied by site, depth, and transect.**

**S2 Table. PERMANCOVA showing that sediment total and available (EDTA extractable) PTEs varied by site and transect.**

**S3 Table. Analysis of Variance for sediment properties. F-statistics of a two-way ANOVA with ‘site’ and ‘depth’ as the two factors. The last two columns indicate whether reference sites; Tyee Banks (TB), Wolfe Cove (WC) and Inverness Passage (IP) are significantly (p < 0.05) different from potentially contaminated sites (Cassiar Cannery and Papermill Bay).**

**S4 Table. Concentrations (mg kg**^**-1**^**) of Cr, Co, Ni, Cu, Zn, As, Cd and Pb in benthic invertebrates sampled from five intertidal mudflats along the north coast of British Columbia, Canada. CC: Cassiar Cannery. WC: Wolfe Cove. IP: Inverness Passage. PB: Papermill Bay. TB: Tyee Banks**

