## Supporting information for "Survey of intertidal ecosystem reveals a legacy of potentially toxic elements from industrial activity in the Skeena Estuary, British Columbia, Canada"

**Quality control**

Total organic carbon and nitrogen

Analysis of sediment samples for total organic carbon and nitrogen content using a Thermo Scientific Flash 2000 Organic Elemental Analyser was conducted alongside one blank every 20 samples and a 5 mg in-house reference soil (107% ± 10.08% recovery for N and 101% ± 0.99% recovery for C, n = 4) traceable to GBW07412, certified for N by State Bureau of Technical Supervision, The People’s Republic of China and to AR-4016, certified for C by Alpha Resources Inc.. The instrument was calibrated with 1 and 3 mg samples of an aspartic acid standard.

ICP-MS analysis of sediment digests

Each batch of 33 samples was digested alongside four blank tubes, two samples of an in-house reference soil (As: 110% ± 6.58%, Cd: 95% ± 4.01%, Co: 81% ± 18.24%, Cr: 91% ± 5.33%, Cu: 88% ± 6.16%, Ni: 79% ± 2.90%, Pb: 97% ± 8.98%, Zn: 104% ± 10.91%, n = 6) traceable to BCR-143R trace elements in a sewage sludge amended soil, certified by Commission of the European Communities, Community Bureau of Reference, and one sample of BCR - 320R Channel Sediment, certified by Commission of the European Communities, Community Bureau of Reference (As: 104% ± 2.44%, Cd: 100% ± 3.16%, Co: 87% ± 3.55%, Cr: 46% ± 2.39%, Cu: 94% ± 4.49%, Ni: 81% ± 3.45%, Pb: 102% ± 4.04%, Zn: 98% ± 2.85%, n = 3). The BCR-320R Channel Sediment is certified for total concentrations, whereas the method adopted here (U.S. EPA Method 3051A) is not intended to accomplish total decomposition of the sample.

Mercury analysis of sediments

The sediments were measured alongside three samples of BCR-320R Channel Sediment, certified by the Commission of the European Communities, Community Bureau of Reference (92% ± 3.03, n = 3) and nine samples of DORM-4 Dogfish Muscle, certified by the National Research Council, Canada (92% ± 2.32, n = 9).

ICP-MS analysis of invertebrate digests

The samples were analysed in two batches, each alongside six blank tubes and two samples of ERM-CE278 Mussel tissue; Commission of the European Communities, Community Bureau of Reference (As: 104% ± 11.10%, Cd: 90% ± 4.78%, Cr: 114% ± 5.74%, Cu: 77% ± 13.39%, Pb: 106% ± 5.54%, Zn: 83% ± 10.19%, n = 4).

**Statistical analysis**

Effect of site and sediment depth on sediment properties and PTE concentrations

The influence of site and sediment depth on sediment PTE concentrations and properties was quantified using two-way analysis of variance using the 18^th^ Edition of Genstat. Normality and homoscedasticity were assessed by inspecting the residual plots and logarithmic or reciprocal transformations made, where necessary. Multiple comparisons were made using the Fisher Least Significant Differences test at the 95% level. Spatial variation of sediment properties (C, N, pH, and median diameter) was assessed using permutational multivariate analysis of covariance, PERMANCOVA (Anderson et al. 2008; Gerwing et al. 2016a). The response variable for the PERMANCOVA was a resemblance matrix calculated using Euclidian distances, and sediment properties were normalized prior to analysis. Before normalization, all variables were square root transformed to correct for skewed distributions (Clarke 1993; Clarke et al. 2008). In the PERMANOVA, site (five levels) was a fixed factor and transect nested within site (5 levels) was a random factor. Depth was included as a covariate. The lowest level of replication was an individual sediment core, n = 76 seines. As part of the PERMANCOVA, we quantified variance components, the proportion of the multivariate variation accounted for by each variable (Searle et al. 1992; Anderson et al. 2008).

The relationship between PTE concentrations and sediment properties

The relationship between PTE concentrations and sediment properties was tested in a step-wise manner. First, using the program PRIMER (Clarke & Gorley 2015), the relationship at all sediment depths, was quantified using PRIMER’s RELATE function. This function compares two resemblance matrices looking for any relationships. In this case, the resemblance matrices were composed of sediment physicochemical property data and sediment PTE data. The sediment physicochemical property resemblance matrix was composed of four variables (pH, %N, %C, and median particle size), all of which were fourth root transformed to correct for skewed distributions. Sediment physicochemical property data were then normalized and a resemblance matrix was constructed from these data using Euclidian distances. The sediment PTE resemblance matrix was constructed in the same manner, except data did not require a transformation to correct for skewed distributions, but data were normalized. As a significant relationship was observed between the two resemblances matrices, this relationship was further explored using Principal Component Analyses on the variance-covariance matrix of all sediment PTE and sediment property data using the 18^th^ edition of Genstat. A PERMANCOVA, conducted as described above, was also used to test the spatial variation in total and extractable PTE concentrations in sediment (n = 98).

The relationship between PTE concentrations in sediments and invertebrates

Relationships between the PTE concentrations observed in the sediments and in the collected invertebrates were examined in several configurations using RELATE, and in all cases the sediment PTE resemblance matrix was constructed as described above on normalized data. First, a sediment PTE resemblance matrix was compared to the observed concentrations of all metals identified in invertebrates, (total and extractable), as well as total and extractable metals analyzed separately. Finally, as not all invertebrates were observed at all sites, we looked for a relationship between contamination (all PTEs, total, and extractable) in two invertebrates observed at each site (*M. balthica*, and *G. oregonensis*) and PTEs in the sediment. For all the analyses described above, all depth levels were retained in the resemblance matrices. When relationships between sediment and invertebrate contaminations were separated by depths, similar relationships were observed. As such, we only present relationships that include all depths here. Non-metric multidimensional (nMDS) scaling plots were used to assess relationships between invertebrates and PTE concentrations. The response variable for all nMDS plots were the resemblance matrices described above, and nMDS plot construction incorporated 100 restarts. All nMDS graphs had a stress ~0.2, and were considered good 2-dimensonal representation of higher dimensional trends (Clarke 1993).

**S1 Table. PERMANCOVA showing sediment properties (pH, median particle diameter, C and N) varied by site, depth, and transect.**

| Source | df | MS | Pseudo-F | Unique Permutations | *p* | Variance Components (%) |
| --- | --- | --- | --- | --- | --- | --- |
| Depth | 1 | 2.23 | 6.44 | 9943 | 0.0006 | 1.52 |
| Site | 4 | 13.17 | 9.15 | 9935 | 0.0001 | 50.24 |
| Transect (site) | 16 | 1.66 | 5.51 | 9887 | 0.0001 | 22.41 |
| Depth X Site | 3 | 1.28 | 4.25 | 9954 | 0.0004 | 3.35 |
| Depth X Transect (Site) | 16 | 0.53 | 1.76 | 9899 | 0.02 | 3.97 |
| Residual | 35 | 0.30 |  |  |  | 18.52 |
| Total | 75 |  |  |  |  |  |

**S2 Table. PERMANCOVA showing that sediment total and available (EDTA extractable) PTEs varied by site and transect.**

| Source | df | MS | Pseudo-F | Unique Permutations | *p* | Variance Components (%) |
| --- | --- | --- | --- | --- | --- | --- |
| Depth | 1 | 572.5 | 1.48 | 9945 | 0.23 | 0.18 |
| Site | 4 | 11480.0 | 14.62 | 9945 | 0.0001 | 50.88 |
| Transect(Site) | 20 | 781.0 | 2.04 | 9902 | 0.002 | 9.50 |
| Depth X Site | 4 | 409.8 | 1.07 | 9937 | 0.39 | 0.13 |
| Depth X Transect(Site) | 20 | 535.1 | 1.40 | 9906 | 0.13 | 3.71 |
| Residual | 48 | 382.0 |  |  |  | 35.60 |
| Total | 97 |  |  |  |  |  |

**S3 Table. Analysis of Variance for sediment properties. F-statistics of a two-way ANOVA with ‘site’ and ‘depth’ as the two factors. The last two columns indicate whether reference sites; Tyee Banks (TB), Wolfe Cove (WC) and Inverness Passage (IP) are significantly (p < 0.05) different from potentially contaminated sites (Cassiar Cannery and Papermill Bay).**

| Variable |  | Site | Depth | Site × Depth interaction | Cassiar Cannery | Papermill Bay |
| --- | --- | --- | --- | --- | --- | --- |
| Sediment pH |  | 24.96*** | 1.39 | 1.27 | IP, TB | WC, TB |
| Median sediment diameter | # | 53.92*** | 0.85 | 1.97* | TB | TB |
| Sediment %C | $ | 49.82*** | 3.67* | 1.35 | TB | WC, TB |
| Sediment %N | $ | 41.00*** | 1.67 | 0.53 | TB | WC, TB |
| Total sediment As |  | 183.99*** | 2.54 | 1.1 | IP | WC, TB |
| Total sediment Cd | $ | 11.49*** | 1.28 | 0.8 | TB | TB |
| Total sediment Co |  | 76.88*** | 2.09 | 1.93* | IP, TB | WC, TB |
| Total sediment Cr | $ | 75.73*** | 0.17 | 0.73 | TB | WC, TB |
| Total sediment Cu |  | 61.51*** | 3.45* | 0.66 | TB | WC, TB |
| Total sediment Hg | # | 41.91*** | 0.13 | 0.77 | TB | WC, TB |
| Total sediment Ni |  | 108.17*** | 1.36 | 0.5 | IP, TB | WC, TB |
| Total sediment Pb | $ | 89.15*** | 0.43 | 0.53 | TB | WC, TB |
| Total sediment Zn |  | 14.27*** | 1.47 | 0.79 | TB | TB |
| EDTA extractable sediment Cd | $ | 8.19*** | 2.62 | 0.47 | TB | WC, TB |
| EDTA extractable sediment Co | $ | 17.72*** | 4.45** | 1.63 | IP | WC |
| EDTA extractable sediment Cr | # | 72.22*** | 11.22*** | 2.67** | TB | WC, TB |
| EDTA extractable sediment Cu |  | 45.67*** | 5.43** | 1.31 | IP, TB | TB |
| EDTA extractable sediment Ni |  | 21.64*** | 10.24*** | 1.86 | IP | WC, TB |
| EDTA extractable sediment Pb |  | 22.97*** | 0.01 | 0.5 | IP, TB | WC, TB |
| EDTA extractable sediment Zn |  | 20.63*** | 9.67*** | 1.31 | TB | WC, TB |

\# = Log_10_ transformed, $ = Reciprocal transformed, * = P< 0.05, ** = P< 0.01, *** = P< 0.001

**S4 Table. Concentrations (mg kg^-1^) of Cr, Co, Ni, Cu, Zn, As, Cd and Pb in benthic invertebrates sampled from five intertidal mudflats along the north coast of British Columbia, Canada. CC: Cassiar Cannery. WC: Wolfe Cove. IP: Inverness Passage. PB: Papermill Bay. TB: Tyee Banks**

| **Site** | **Common name** | **Latin name** | **As** | **Cd** | **Co** | **Cr** | **Cu** | **Ni** | **Pb** | **Zn** |
| --- | --- | --- | --- | --- | --- | --- | --- | --- | --- | --- |
| CC | Clams | *Macoma balthica* | 6.25 | 0.05 | 1.67 | 3.72 | 20.59 | 3.38 | 0.90 | 39.92 |
| IP | Clams | *Macoma balthica* | 4.59 | 0.07 | 1.76 | 3.69 | 21.11 | 2.39 | 0.95 | 30.42 |
| TB | Clams | *Macoma balthica* | 7.59 | 0.07 | 1.24 | 2.17 | 20.80 | 1.50 | 0.55 | 29.24 |
| WC | Clams | *Macoma balthica* | 3.19 | 0.04 | 1.49 | 3.76 | 34.60 | 1.99 | 0.70 | 30.73 |
| PB | Clams | *Macoma balthica* | 2.09 | 0.03 | 0.71 | 1.14 | 44.49 | 0.98 | 0.27 | 16.62 |
| CC | Blue mussels | *Mytilus edulis* | 2.16 | 0.44 | 0.89 | 1.45 | 3.44 | 0.98 | 0.23 | 8.01 |
| WC | Blue mussels | *Mytilus edulis* | 5.00 | 0.82 | 0.66 | 0.99 | 3.71 | 1.01 | 0.53 | 18.13 |
| IP | Blue mussels | *Mytilus edulis* | 6.43 | 1.26 | 3.79 | 4.60 | 9.50 | 3.10 | 1.04 | 27.75 |
| CC | Horse mussels | *Modiolus rectus* | 1.74 | 0.29 | 0.88 | 2.30 | 5.67 | 1.36 | 0.67 | 6.91 |
| WC | Soft shell clams | *Mya arenaria* | 4.22 | 0.20 | 2.72 | 7.43 | 9.36 | 3.21 | 1.11 | 21.14 |
| PB | Soft shell clams | *Mya arenaria* | 2.43 | 0.12 | 1.05 | 1.48 | 3.70 | 0.87 | 0.72 | 13.96 |
| CC | Soft shell clams | *Mya arenaria* | 6.80 | 0.22 | 2.21 | 4.29 | 8.26 | 3.49 | 1.01 | 18.38 |
| IP | Soft shell clams | *Mya arenaria* | 15.61 | 0.39 | 6.41 | 6.41 | 13.36 | 4.93 | 1.83 | 47.16 |
| WC | Bent nose macoma | *Macoma naustia* | 3.82 | 0.11 | 2.78 | 6.82 | 22.97 | 3.64 | 1.00 | 30.52 |
| WC | Bent nose macoma | *Macoma naustia* | 5.98 | 0.19 | 4.45 | 10.38 | 42.02 | 4.96 | 1.49 | 57.06 |
| CC | Green shore crabs | *Hemigrapsus oregonensis* | 7.28 | 0.25 | 1.76 | 3.15 | 82.84 | 2.45 | 0.76 | 53.81 |
| IP | Green shore crabs | *Hemigrapsus oregonensis* | 8.91 | 0.34 | 1.68 | 2.59 | 116.89 | 2.19 | 0.61 | 88.73 |
| WC | Green shore crabs | *Hemigrapsus oregonensis* | 6.05 | 0.27 | 0.71 | 1.33 | 74.81 | 1.17 | 0.76 | 54.80 |
| WC | Hermit crabs | *Pagurus hirsutiusculus* | 5.55 | 0.21 | 1.46 | 3.99 | 100.06 | 2.59 | 1.31 | 51.29 |
| CC | Hermit crabs | *Pagurus hirsutiusculus* | 4.01 | 0.26 | 0.34 | 0.48 | 71.57 | 0.38 | 0.14 | 50.78 |
| CC | Isopods | *Gnorimosphaeroma oregonensis* | 4.52 | 0.41 | 1.68 | 1.91 | 125.23 | 1.59 | 0.43 | 43.51 |
| TB | Isopods | *Gnorimosphaeroma oregonensis* | 5.55 | 0.67 | 2.61 | 2.82 | 186.78 | 2.30 | 0.64 | 50.52 |
| WC | Isopods | *Gnorimosphaeroma oregonensis* | 3.37 | 0.20 | 1.11 | 2.23 | 88.58 | 1.23 | 0.34 | 47.07 |
| IP | Isopods | *Gnorimosphaeroma oregonensis* | 5.47 | 0.46 | 1.58 | 1.48 | 130.14 | 1.33 | 0.48 | 63.75 |
| PB | Isopods | *Gnorimosphaeroma oregonensis* | 5.42 | 0.36 | 2.54 | 6.96 | 109.07 | 3.38 | 0.81 | 87.51 |
| WC | Kelp isopods | *Idotea wosnesenskii* | 6.54 | 0.43 | 1.62 | 2.54 | 47.23 | 2.10 | 0.68 | 63.27 |
| IP | Kelp isopods | *Idotea wosnesenskii* | 8.85 | 0.52 | 1.96 | 1.28 | 60.00 | 1.88 | 0.42 | 56.48 |
| WC | Ghost shrimp | *Neotrypaea californiensis* | 14.30 | 1.32 | 1.79 | 1.89 | 177.56 | 1.16 | 0.62 | 80.09 |
| TB | Amphipods | *Eogammarus confervicolus* | 3.94 | 0.32 | 1.80 | 2.49 | 89.84 | 2.37 | 0.75 | 65.63 |
| CC | Amphipods | *Eogammarus confervicolus* | 2.59 | 1.30 | 1.54 | 2.94 | 118.40 | 4.27 | 5.51 | 52.68 |
| PB | Amphipods | *Eogammarus confervicolus* | 4.06 | 0.39 | 1.34 | 1.38 | 87.68 | 1.00 | 0.75 | 78.42 |
| IP | Amphipods | *Eogammarus confervicolus* | 4.53 | 0.37 | 1.88 | 2.45 | 99.22 | 0.72 | 2.34 | 41.55 |
| IP | Amphipods | *Americorophium salmonis* | 9.99 | 0.51 | 3.57 | 8.16 | 75.63 | 4.79 | 3.17 | 73.33 |
| CC | Amphipods | *Americorophium salmonis* | 8.39 | 0.65 | 2.96 | 7.56 | 111.65 | 5.21 | 5.04 | 202.87 |
| PB | Amphipods | *Americorophium salmonis* | 5.78 | 0.64 | 1.62 | 5.07 | 126.15 | 1.57 | 4.43 | 87.00 |
| CC | Lugworms | *Abarenicola pacifica* | 62.63 | 0.34 | 12.99 | 16.68 | 28.24 | 26.21 | 3.93 | 88.45 |
| PB | Lugworms | *Abarenicola pacifica* | 23.87 | 0.32 | 11.63 | 23.57 | 26.77 | 19.32 | 4.06 | 105.21 |
| WC | Lugworms | *Abarenicola pacifica* | 13.69 | 0.24 | 9.09 | 25.06 | 28.92 | 16.98 | 4.32 | 75.11 |
| WC | Goniadidae worms | *Glycinde picta* | 13.35 | 0.99 | 1.04 | 0.91 | 9.08 | 2.98 | 0.27 | 202.17 |
| WC | Light edged ribbon worm | *Cerebratulus californiensis* | 7.82 | 2.56 | 0.34 | 0.31 | 39.14 | 0.07 | 0.16 | 101.17 |
| CC | Purple Backed Ribbon worms | *Paranemertes peregrina* | 25.36 | 1.13 | 3.81 | 0.78 | 23.18 | 1.51 | 5.64 | 201.74 |
| WC | Purple Backed Ribbon worms | *Paranemertes peregrina* | 8.94 | 1.15 | 3.50 | 10.06 | 23.23 | 6.15 | 2.77 | 134.76 |
| WC | Catworm | *Nephtys caeca* | 22.62 | 4.08 | 2.30 | 2.27 | 10.30 | 3.64 | 29.61 | 142.64 |
| CC | Catworm | *Nephtys caeca* | 12.59 | 6.57 | 2.94 | 1.63 | 30.95 | 3.25 | 40.96 | 180.33 |
| CC | Clam worm | *Nereis vexillosa* | 11.22 | 5.34 | 4.09 | 2.32 | 23.09 | 2.21 | 38.28 | 341.61 |
| WC | Paraonidae worms | *Aricidea hartleyi* | 8.31 | 1.08 | 14.36 | 52.78 | 47.39 | 21.76 | 10.71 | 106.83 |
| CC | Spionidae tube worms | *Streblospio benedicti* | 22.57 | 6.21 | 6.02 | 15.91 | 24.65 | 10.37 | 7.00 | 185.84 |
| WC | Sandworm | *Alitta brandti* | 11.71 | 1.02 | 4.46 | 15.13 | 23.31 | 6.62 | 10.74 | 126.03 |
